# Evaluation of MeaSeq: comprehensive analysis and reporting of measles virus whole genome sequences

**DOI:** 10.64898/2026.05.12.724559

**Authors:** Darian Hole, Ahmed Abdalla, Vanessa Zubach, Molly Pratt, Stephanie Van Driel, Samar Ashfaq, Joanne Hiebert, Ana T. Duggan

## Abstract

Although vaccine-preventable, measles virus (MeV) continues to pose a significant public health challenge, with a substantial resurgence of cases worldwide. As whole-genome sequencing (WGS) becomes increasingly affordable and routinely adopted in public health laboratories, reliable and accessible analysis of next-generation sequencing (NGS) data is critical for outbreak investigation and molecular surveillance. Here, we present MeaSeq, a fast, user-friendly, open-source bioinformatics pipeline for MeV analysis using Illumina or Oxford Nanopore Technologies (ONT) NGS data. MeaSeq performs quality control assessments, consensus genome assembly and variant detection, optional genotype-specific reference selection, Distinct Sequence Identifier (DSId) assignment via user-provided databases or hashing, sub-consensus variant visualization, genome quality assessment, and standardized HTML reporting. We compared the performance of MeaSeq on NGS data generated from multiple sequencing platforms and targeted enrichment strategies against gold-standard Sanger data, reference genomes, and publicly available comparative data. This validation demonstrates that MeaSeq provides an accurate, reproducible, and accessible solution for routine MeV WGS analysis, supporting genomic surveillance and outbreak response workflows in public health and research settings.

**Impact Statement:** The recent surge in measles cases worldwide, causing several countries to lose their measles elimination status, underscores the urgent need for effective and accessible genomic surveillance. Our manuscript introduces MeaSeq, a comprehensive and open-source bioinformatics pipeline specifically designed for analyzing MeV NGS data. MeaSeq includes MeV specific analyses such as genotype prediction from sequencing reads with optional genotype-specific reference selection; DSId assignment; quality control checks such as genome rule-of-six divisibility and gene CDS validation; subconsensus nucleotide analysis with mixed-site highlighting; and genomic plotting. By leveraging NGS technology, our pipeline can facilitate the identification of transmission chains and may provide critical insights into the dynamics of MeV outbreaks. This information is essential for public health officials and researchers to implement targeted interventions and optimize vaccine strategies. Additionally, the open-source nature of MeaSeq fosters collaboration and innovation within the scientific measles community along with providing access to a wider range of researchers.

**Data Summary:** The MeaSeq pipeline code is available on GitHub (https://github.com/phac-nml/measeq). Comparative datasets of publicly available WGS data were accessed through the NCBI Sequence Read Archive under the following BioProjects:

PRJNA869081 (https://www.ncbi.nlm.nih.gov/bioproject/PRJNA869081)

PRJNA480551 (https://www.ncbi.nlm.nih.gov/bioproject/PRJNA480551)

PRJNA1017431 (https://www.ncbi.nlm.nih.gov/bioproject/PRJNA1017431)

PRJNA1241325 (https://www.ncbi.nlm.nih.gov/bioproject/PRJNA1241325)

PRJNA1174053 (https://www.ncbi.nlm.nih.gov/bioproject/PRJNA1174053)

PRJNA1293457 (https://www.ncbi.nlm.nih.gov/bioproject/PRJNA1293457)

PRJNA843031 (https://www.ncbi.nlm.nih.gov/bioproject/PRJNA843031)

Whole-genome sequences were included in the validation analysis if they consisted of paired-end data (Illumina) and achieved ≥95% genome completeness following trimming of the 5′ and 3′ untranslated regions (UTRs). This criterion ensured sufficient genome coverage for robust validation while allowing for limited missing data arising from regions of low sequencing depth or amplicon dropout.

A complete list of sequences included in the validation, along with their accession numbers, is provided in Supplementary Table 1.

## Introduction

Measles virus (MeV) is one of the most transmissible viruses known to humans [1]. Common symptoms of measles include fever, cough, runny nose, conjunctivitis, and rash [2]. Approximately 1 to 3 per 1000 people infected with MeV die from respiratory or neurologic complications [2]. Measles disease can be prevented with the safe and highly effective live-attenuated vaccine.

Measles is a negative strand RNA virus belonging to the genus Morbillivirus within the Paramyxoviridae family [3]. These viruses must conform to the rule-of-six to comply with replication competency [3], meaning that their genome length must be divisible by six. Insertions and deletions (indels) have been reported [4, 5], some resulting in non-standard genome lengths, but they always comply with the rule-of-six. Multiple studies suggest that these indels are predominantly localized to the non-coding region between the matrix and fusion genes (MF-NCR) [6, 7].

The World Health Organization (WHO) recommends molecular surveillance of every MeV outbreak by genotyping the 450 nucleotide (nt) region of the C-terminus end of the nucleoprotein (N) gene (N450) [8]. Distinct Sequence Identifiers (DSIds) are assigned to each unique N450 sequence that is submitted to the global WHO measles database (MeaNS2 https://who-gmrln.org/means2). The use of DSIds can aid in epidemiological surveillance and tracking purposes [9]. Historically, there were 24 MeV genotypes circulating globally, but as of 2021 only 2 genotypes (D8 and B3) remain active [10]. Due to this decreasing genetic diversity, genotyping the N450 is not always adequate for monitoring transmission, and a higher resolution of sequencing is needed to resolve outbreaks [9]. Additional sequencing of the hypervariable, noncoding region between the matrix (M) and fusion (F) genes (MF-NCR) and/or whole genome sequencing (WGS) has been suggested to further refine the monitoring of transmission chains, and an increasing number of laboratories have already successfully used this approach [11].

Recently, MeV activity has drastically increased in several countries, resulting in large or disruptive outbreaks [12, 13]. In 2025, after 27 years of measles elimination status, the Pan American Health Organization (PAHO) revoked Canada’s measles elimination status due to a large multi-jurisdictional outbreak of a measles strain that had been circulating longer than 12 months [14]. In 2026, the United Kingdom, Spain and several other European countries have also lost their measles elimination status [15]. The resurgence, transmissibility, and decreasing diversity of measles in many parts of the world highlights the need for effective surveillance and vigorous outbreak tracing. Next generation sequencing (NGS) has become a highly valuable, more affordable, and widely utilized tool for outbreak resolution as shown with its effectiveness during the SARS-CoV-2 pandemic. However, the analysis of NGS data can be costly or challenging for non-bioinformaticians to generate accurate consensus sequences and the required surveillance reporting metrics. Rigorous primer trimming and post-NGS quality control are essential for WGS consensus generation, as failure to remove primer-derived bases and low-quality regions can lead to erroneous consensus sequences and compromise molecular surveillance and outbreak investigations.

Many bioinformatic pipelines have been developed to automate the analyses of NGS data generically from viral samples, such as VirPipe [16], GenomeDetective [17], nf-core/viralrecon [18], and Illumina VSP Pipeline [19]. Initial evaluations of those pipelines showed promising results but did not fully meet the operational requirements for MeV surveillance. Some viral pathogens have customized bioinformatic pipelines to meet their virus-specific needs such as RSV-GENOScan for respiratory syncytial virus analysis [20], COVIDSeq for SARS-CoV-2 [21], and nf-flu for Influenza A and B [22].

Currently, there are no validated pipelines specific for the genomic analyses of MeV. Thus, with decreasing diversity confounding the current genotyping method along with specific genomic constraints and the current pressing outbreaks, there is a demand for an easy-to-use, automated, robust, and validated tool to analyze and visualize NGS data for MeV.

In this study, we developed and thoroughly validated MeaSeq, a bioinformatics pipeline specific for MeV WGS data generated using both hybrid capture or tiled amplicon methods with either Illumina or Oxford Nanopore Technologies (ONT) platforms. MeaSeq was created to expand upon all-inclusive viral pipelines including nf-core/viralrecon and the ARTIC nanopore pipeline [23] to apply MeV-specific parameters and quality checks with a focus on final reporting metrics and plots. The design was informed by operational requirements within the Public Health Agency of Canada (PHAC), particularly rapid turnaround times for outbreak response, standardized reporting outputs for epidemiological use, the integration of the DSId system, and the need to support users with varying levels of bioinformatics expertise. MeaSeq accomplishes MeV-specific requirements through several sub-analyses including genotype prediction and optional genotype specific reference analysis, DSId assignment, rule-of-six genome divisibility checks, validation of intact gene CDS regions using Nextclade [24], subconsensus variation analysis, and genomic depth and quality plotting. All of which are summarized in a final easy to use standalone interactive HTML report.

Our study evaluated MeaSeq’s ability and accuracy in generating N450 surveillance and whole-genome consensus sequences. The performance was first validated with gold-standard Sanger-generated N450 and MF-NCR sequences in addition to ATCC MeV reference genomes for genotypes D8 and B3. MeV whole-genome consensus sequences from NCBI, for which both GenBank consensus sequences and raw sequencing reads (SRA) were available, were compared with the MeaSeq generated consensus sequences using the same raw data. This comparison allowed for the assessment of a diverse range of MeV strains and library qualities.

## Materials and Methods

### MeaSeq pipeline

MeaSeq is a command line pipeline for the analysis of MeV NGS data obtained using hybrid capture or tiled amplicon methods on Illumina or ONT platforms. It is designed for rapid outbreak response and routine surveillance producing consensus genomes, variant calls, and detailed sample level QC (Figure 1). MeaSeq is written in Nextflow [25] following community standards [26] and can be run with a multitude of containerized approaches ensuring its reproducibility and availability across a variety of computational platforms.

**Figure 1.**
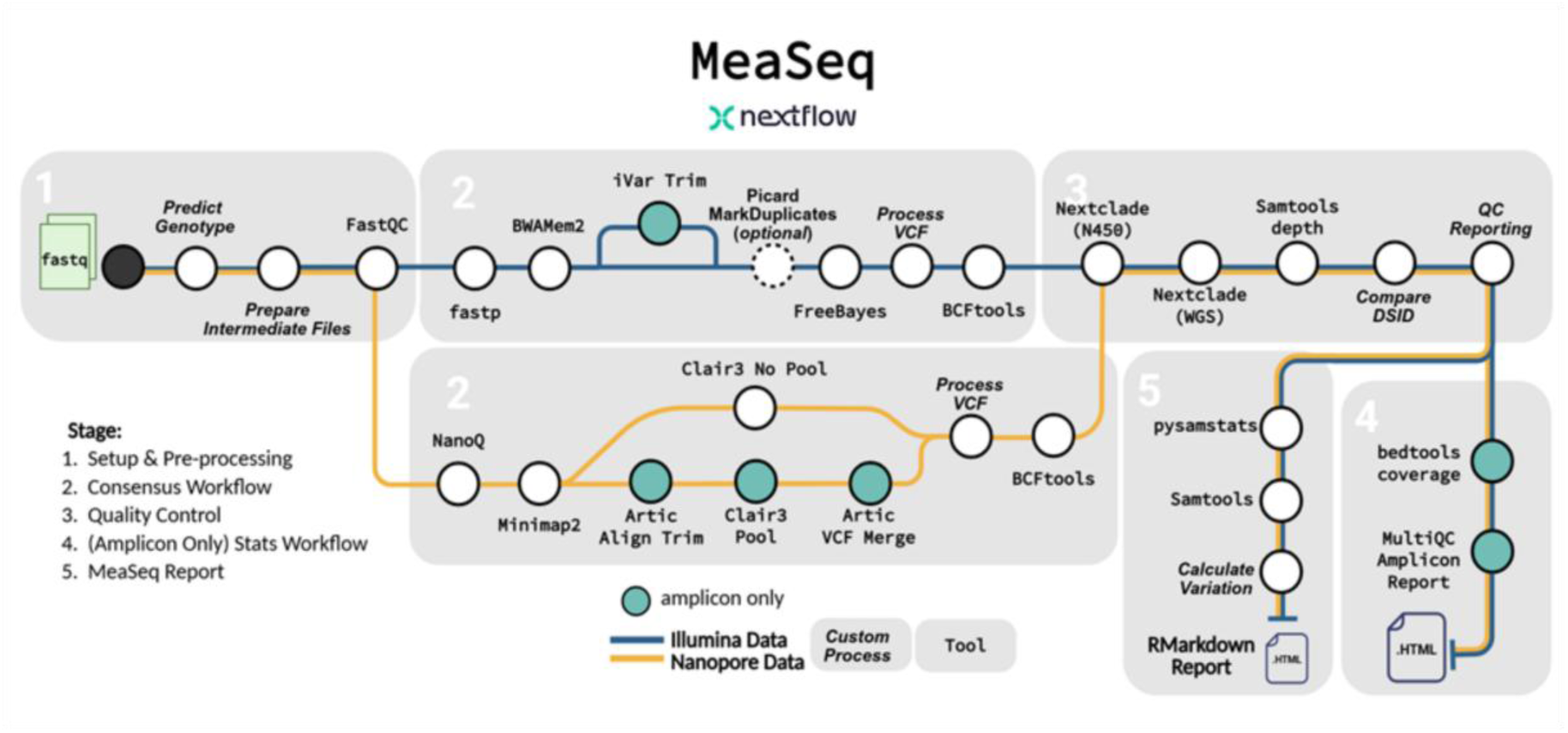
MeaSeq v1.1.0 workflow. MeaSeq performs quality assessment and generates MeV consensus sequences from either Illumina (blue) or Nanopore (orange) reads. Amplicon-specific processing steps are indicated by the blue-filled circles. The final comprehensive RMarkdown report summarizes the generated consensus sequences and any detected variation compared to genotypic reference sequences. Figure created in BioRender.

### Inputs and Configuration

The MeaSeq pipeline requires the input of single or paired-end FASTQ reads through a CSV formatted sample sheet. Optional metadata can be provided as a separate TSV file containing a sample column that matches the sample names in the sample sheet. During execution, the pipeline joins the metadata and genomic analysis results into final summary tables. At minimum, the path to the sample sheet, the directory where results should be saved, and a specification for Illumina or ONT data is required to run the pipeline.

### Reference Genotype Assignment

A central design element of MeaSeq is the assignment of each sample to the most appropriate MeV genotype before assembly. MeaSeq implements a genotype classification module that compares sequencing reads against a supplemented WHO N450 reference dataset. This dataset is used to distinguish circulating MeV genotypes based on the N450 sequence. The output of this prediction step is used to determine which reference genome and primer scheme (if running amplicon data) should be used for read mapping, consensus assembly, and final QC.

MeaSeq is currently preset with a reference genome and primer BED file for three measles genotypes with support for all 24 measles genotypes available. The presets encompass the two currently circulating genotypes (B3 and D8) and the vaccine genotype (A). Users modify and customize the genotype reference FASTA and primer BED files using a parameters config file or with command line arguments.

Genotype specific processing can be disabled by specifying a single reference sequence for all samples. These options allow flexibility for users to extend the pipeline to emerging genotypes or utilize new primer schemes without modifying the underlying code.

### Illumina Consensus Workflow

Following genotype assignment and the generation of read-level quality reports by FASTQC [27], the MeaSeq workflow diverges based on the specified sequencing platform (Figure 1). For Illumina read processing, Fastp is used for adapter and quality trimming [28]. Trimmed reads are then aligned to the designated reference using BWA-Mem2 [29]. For amplicon libraries, primer sequences are removed using iVar [30]. Illumina optical duplicate reads can be marked and removed with Picard MarkDuplicates [31]. This option is disabled by default in the interest of maintaining read depth in tiled amplicon designs, while providing flexibility for non-amplicon-based libraries where duplicates may represent technical artifacts. Candidate variant calling is performed using Freebayes [32] with final variants and ambiguous positions determined with a set of filtering parameters. Initially, candidate variants are filtered based on a minimum quality score of 20, set based on the Freebayes documentation, along with a minimum depth of 10 reads. Variants with allele frequencies between 0.30 and 0.75 are considered ambiguous due to discordance in the underlying sequencing data which may stem from viral evolution or underlying sequencing artifacts. Variants with an allele frequency greater than 0.75 are considered to pass and are designated as a single nucleotide polymorphism (SNP) for that position. Indel variants are applied at a lower passing frequency of 0.60 to account for alignment difficulties in the lower complexity regions of the MeV genome. These parameters can be adjusted using a parameters config file or on the command line; the default values were determined from comparisons to the Sanger validation dataset in addition to using previous SARS-CoV-2 analyses as a starting point [33]. Finally, the consensus sequence is generated by BCFtools [34] incorporating detected variants and masking regions below the minimum defined depth of 10 reads with Ns. See Table 1 for a list of tools used by MeaSeq and their default values.

**Table 1.** MeaSeq Default Parameters.

| Parameter Description | Tool | Default Arguments | Short Summary |
| --- | --- | --- | --- |
| Read filtering | Illumina: fastp | --cut_front --cut_tail --trim_poly_x --cut_mean_quality 25 --average_qual 25 --length_required 100 | Trimming low quality positions on the starts and ends of reads along with filtering out overall poor-quality reads and reads that are shorter than 100bp |
|  | Nanopore: nanoq | -l 200 -q 7 | Filtering reads that are shorter than 200bp and have an overall quality score less than 7 |
| Alignment | Illumina: bwamem2 mem |  | No additional optional arguments to the bwamem2 aligner |
|  | Nanopore: Minimap2 | -a -x map-ont | Output the alignment in SAM format with the map-ont default parameters set |
| Alignment filtering | Illumina: samtools view | -b -f 3 -F 2048 | Include only properly paired reads and remove supplementary alignments |
|  | Nanopore: samtools view | -F 2308 | Remove supplementary alignments and unmapped reads |
| Primertrimming | Illumina: iVar trim | -x 0 | Trim primers based on position with no additional offset by default |
|  | Nanopore: artic align_trim | --trim-primers --primer-match-threshold 15 | Trim primers found based on position and primer pool with a default match threshold of 15 |
| Minimum depth | Illumina: make_depth_mask | --depth 10 | Mask sites that are below the minimum depth of 10 with an N |
|  | Nanopore: make_depth_mask | --depth 10 | Mask sites that are below the minimum depth of 10 with an N |
| Variant calling | Illumina: freebayes | -p 1 -C 1 -F 0.05 --pooled-continuous --min-coverage 5 --standard-filter | Call potential variants using a ploidy of 1, a minimum candidate allele frequency of 0.05, and standard filters of MapQ 30 and BaseQ 20 outputting all variants called to be further filtered |
|  | Nanopore: clair3 | --haploid_sensitive --enable_long_indel --include_all_ctgs --chunk_size=5000 --ref_pct_full=1 --var_pct_full=1 --no_phasing_for_fa --enable_variant_calling_at_sequence_head_and_tail | Haploid sensitive variant calling using the users clair3 model with realignment of all variants and reference sites. Variant calling at the start and the ends of reads enabled to properly call variants in amplicon data |
| Consensus variant allele frequency filtering | Illumina: process_vcf | -l 0.25 -u 0.75 -m 0.60 | Variants with an AF greater than -u are applied to the final consensus. Variants between -u and -l are set to an ambiguous base matching the mixture of bases at the site. Minimum indel AF threshold -m is set slightly lower |
|  | Nanopore: vcf_filter | --min-allele-freq 0.60 --min-threshold-depth 0.05 | Variants with an AF greater than the minimum allele frequency are applied to the consensus. Variants with less than an AF of the minimum are masked with an N. Variants calls where less than 5% of the positions overall reads are used to make the call are also skipped |
| Consensus variant quality filtering | Illumina: process_vcf | --min-quality 20 | Minimum freebayes variant quality for variant to be included |
|  | Nanopore: vcf_filter | --min-qual 7 | Minimum variant quality for variant to be included. Variants below a variant quality of 2 are skipped and variants between 2 and the minimum are masked with an N |
| Optical duplicate removal | Illumina: Picard markduplicates | --ASSUME_SORTED true --VALIDATION_STRINGENCY LENIENT --REMOVE_DUPLICATES true | Optional workflow to remove optical duplicates with Picard |
|  | Nanopore: NA |  |  |

### Nanopore Consensus Workflow

For ONT data, NanoQ [35] performs the quality filtering with a minimum read quality of 12. The filtered reads are aligned to the reference genome using Minimap2 [36]. For amplicon data, ARTIC align_trim filters reads that do not correspond with the start and end positions of individual amplicons defined in the BED file [23]. The retained reads are trimmed of terminal bases originating from the primers thereby retaining only target sequences to prevent ambiguity in primer regions and ensure accurate variant calls, see supplementary Figure 1. Candidate variant calls are performed using Clair3 [37] with a variant calling model chosen by the user determined by the basecalling model and flowcell. Additionally, amplicon data filters alignments to their primer pool to call candidate variants to prevent abrupt depth disparities at overlaps which can mask variants. The candidate variants are retained if they meet a minimum depth of 10 and a Phred quality score of ≥7. This corresponds to an approximate 80% relative confidence in the base call being correct based on the variant calling model. A minimum allele frequency threshold of 0.60 is used for both SNPs and indels to allow variants in homopolymer tracts that frequently experience alignment challenges causing lower allele frequencies in these sites. Indels are additionally filtered at a higher Phred quality score of ≥30 to prevent frameshifts and disruptions to the rule-of-six that are not strongly supported. The default values were determined from comparisons to the Sanger validation dataset and ATCC reference genome comparisons. Sites failing any of the variant filters are masked as Ns and applied to the final consensus sequence along with the final passing variants using BCFtools. See Table 1 for a list of tools used by MeaSeq and their default values.

### Nextclade QC, DSId Assignment, and Amplicon Statistics

Following consensus generation, all sequences are evaluated with Nextclade in two steps. First, the N450 region is compared to the Nextclade WHO N450 measles dataset to assign a final genotype to the sample and calculate the completeness of the N450 region. Additionally, a full genome Nextclade run using a custom dataset evaluates gene integrity and flags mutations and frameshifts.

Although DSIds are not publicly available, users have the option to supply previously identified and assigned DSId N450 sequences in a local fasta file. If such a file is provided, MeaSeq compares each sample’s N450 sequence against the DSId reference panel to assign known DSIds. Unmatched sequences or runs where no DSId file is specified will assign a novel identifier based on the MD5 hash of the N450 sequence. This hash remains consistent between runs and therefore allows samples without a known DSId to be preliminarily grouped based on matching MD5 hashes. Samples that have any missing or ambiguous data in the region will be marked as N450 failures as only consensus nucleotides are used to create DSIds.

When tiled amplicon data (--amplicon) is indicated, MeaSeq computes detailed amplicon level depth and completeness metrics. Sequencing depth across each amplicon is summarized using bedtools [38], while custom scripts are used to calculate amplicon completeness. These amplicon statistics are visualized using MultiQC [39] to help identify potential issues in library prep, weak primer pairs, amplicon drops, or issues with the primer files themselves.

### Final Reporting

For each sample, subconsensus analysis is performed on the pileup files using a custom python script to summarize all positions where ≥15% of the reads call a non-reference base. This threshold is lower and more sensitive than the Illumina minimum ambiguous base parameter threshold of 30% to identify potential sequencing or analysis errors and to better track outbreaks by exploring sites with mixed populations which may be the result of current viral evolution. The per-base mean sequencing quality and the proportion of ambiguous nucleotide (‘N’) calls are summarized with pysamstats baseq [40] and samtools [41] to determine regions of low sequencing quality. At this stage, each consensus sequence is checked to ensure that the final genomic length conforms to the rule-of-six. After all checks and quality metrics are computed, the pipeline culminates by summarizing the output in a locally stored HTML report that can be viewed from the user’s web browser. This report displays generic and MeV-specific quality metrics per sample including clear warnings when there is a quality issue detected (Figure 2), and sample specific in-depth stats, variant summaries (Figure 3A), coverage plots and positional quality metrics (Figure 3B). The final reports provide metrics that inform decisions regarding the quality of the sequencing run and each sample individually.

**Figure 2.**
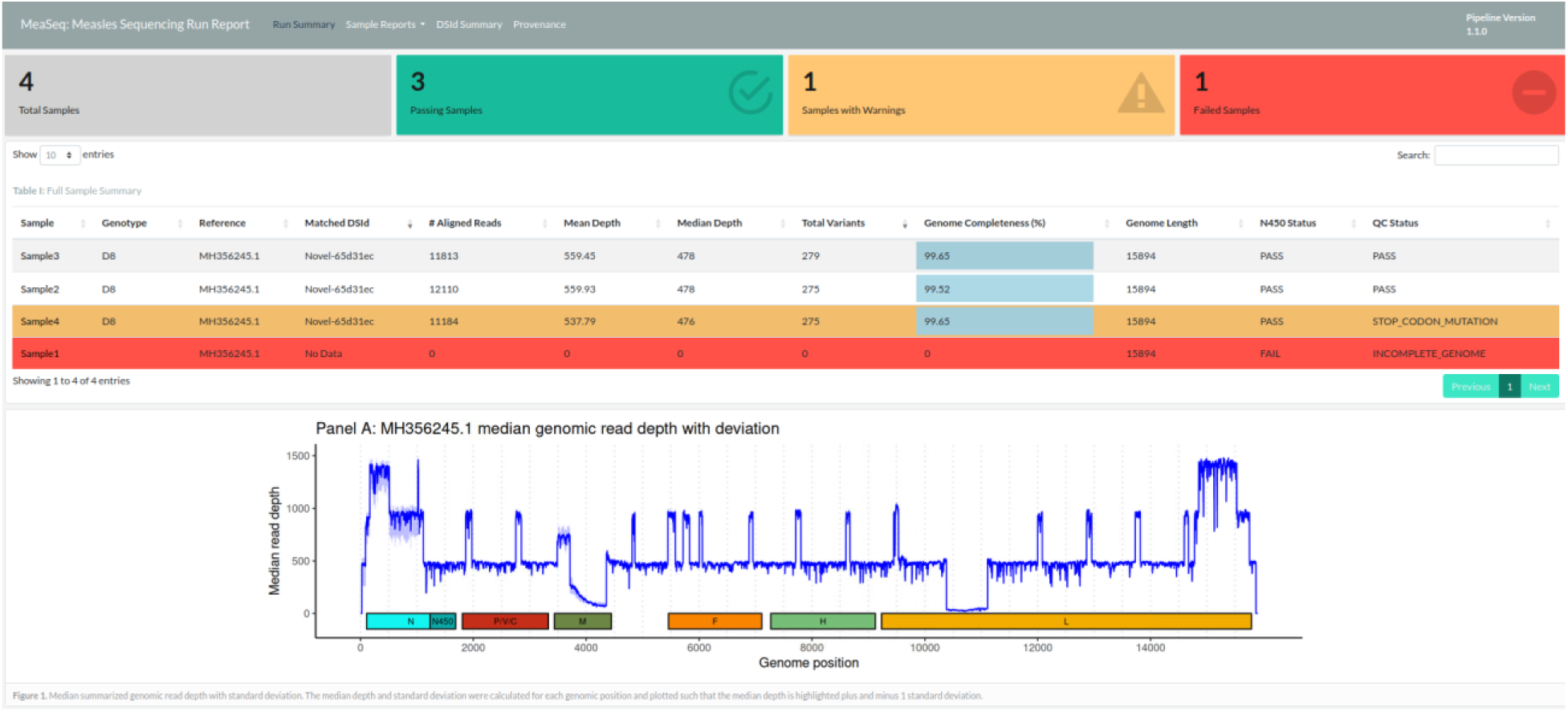
Landing page of MeaSeq report. Summary metrics and MeV-specific warnings for all samples are displayed in the Run Summary tab. For example, a specimen with a late stop codon in the phosphoprotein gene is highlighted in orange. Additional tabs containing sample-specific information, DSId summaries, and tool provenance can be accessed from the report header.

**Figure 3.**
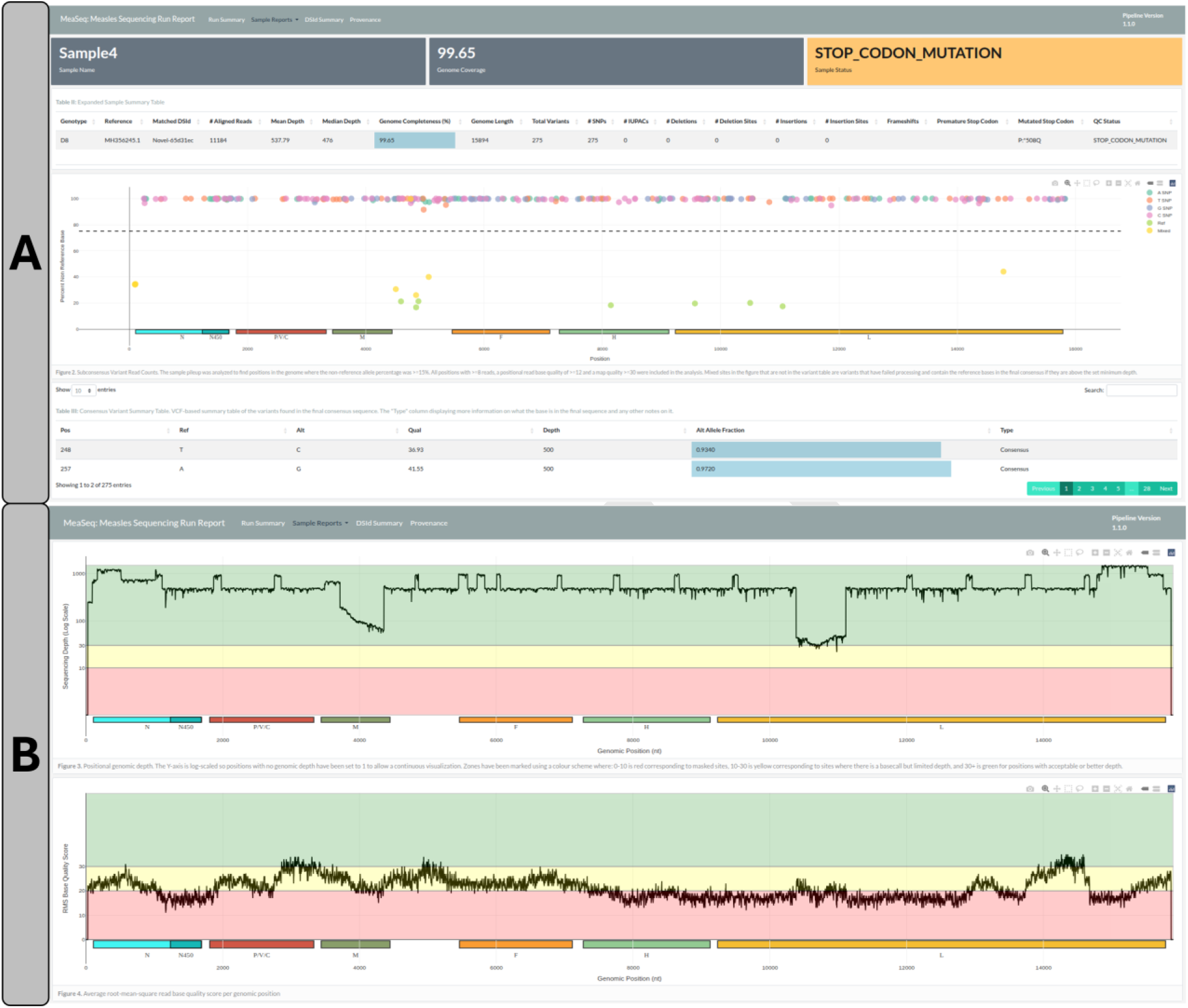
Sample-specific information page of MeaSeq report. The Sample Reports tab includes more in-depth single sample information such as the location of late stop codon in the phosphoprotein gene and variant information (A) as well as interactive per-sample plots with detailed information about sequence and quality variation across each consensus genome (B).

### Gold-standard and reference sequences used for pipeline validation

The validation dataset is comprised of genotype B3 and D8 sequences generated from MeV RT-PCR–positive specimens referred to the PHAC National Microbiology Laboratory (NML) for measles genotyping (n = 146). Sanger sequencing data for the WHO-standardized N450 region and the MF-NCR were used as the gold standard for validation of MeaSeq generated consensus sequences. Sanger amplification and sequencing were performed as previously described [42]. Consensus sequences were generated using SeqMan Pro v11 (DNASTAR Lasergene, WI). Reference-mapped, post-assembly reads were manually trimmed to remove primer-derived read ends, yielding high-quality consensus sequences that were exported in FASTA format. N450 sequences were trimmed to the WHO-standardized window, while MF-NCR sequences were trimmed to include the stop codon of the M open reading frame (ORF) and the start codon of the F ORF, resulting in sequence lengths of either 1,018 or 1,024 nucleotides [42].

For validation, Sanger-derived consensus sequences were compared with NGS data generated using three different WGS approaches: (i) Illumina hybrid capture sequencing, as described by Hiebert *et al.* [42] (BioProject PRJNA1017431; n = 59); (ii) ONT tiled amplicon sequencing, as described by Zubach *et al.* [43] (BioProject PRJNA1174053; n = 36); and (iii) Illumina tiled amplicon sequencing using an NML in-house protocol [44] (n = 51). For the Illumina and ONT tiled amplicon methods, reference MeV virus isolate genomes were obtained from ATCC (ATCC VR-1980 and ATCC VR-1981), sequenced, and the MeaSeq generated output consensus was compared to the ATCC published sequences.

For consistency across analyses, all WGS consensus sequences were trimmed to exclude genomic termini (WGS-t), beginning at the start codon of the N ORF and ending at the stop codon of the L ORF. For most samples, consensus sequences achieved a minimum of 10× coverage across the genome.

Trimmed whole-genome consensus sequences were 15,678 or 15,684 nucleotides in length. A complete list of samples included in the validation set, along with accession numbers, sequencing platform, library preparation method, genotype, and instrument details, is provided in Supplementary Table 1.

### Data Analysis of the validation sequences

The MeaSeq pipeline v1.1.0 was run on all samples using default parameters to generate MeV consensus sequences for comparison to the gold standard methods described above. Primer location BED files were added to the analysis of the amplicon data to properly trim primers and a Clair3 model of “r1041_e82_400bps_sup_v410” was used for the analysis of the ONT data. Final whole genome comparisons of consensus sequences of samples with ≥95% genome completeness were performed using a MAFFT version 7.526 [45] alignment to the genotype specific reference with a custom python script to compare each alignment position and calculate the percent identity between the sequences.

Any samples differing from the validation dataset were further investigated using the MeaSeq report plots and the Integrative Genomics Viewer (IGV) version 2.19.7 [46] to confirm or resolve any genomic sites with allele mismatches or ambiguity using the pileup information. Samples with available Sanger N450 or MF-NCR data were separately aligned to the whole-genome consensus sequences with MAFFT to validate their concordance.

### Collection of publicly available datasets

Publicly available WGS data were identified by screening NCBI BioProjects for use as comparison datasets. BioProjects were included if they contained both WGS-derived consensus sequences and corresponding raw sequencing reads; met a minimum genome completeness threshold of ≥95% after assembly; and, for Illumina datasets, provided properly paired reads. This search yielded data from 223 samples across seven publicly available MeV BioProjects.

Illumina hybrid capture data (n = 107) were obtained from BioProjects PRJNA1017431 [42], PRJNA869081 [47], PRJNA480551 [48], and PRJNA1241325 [49]. Illumina tiled amplicon data (n = 72) were obtained from BioProject PRJNA1293457 [50]. ONT tiled amplicon data (n = 44) were obtained from BioProjects PRJNA1174053 [43] and PRJNA843031 [51]. Details for all included samples, including accession numbers, library preparation method, genotype, and sequencing platform, are provided in Supplementary Table 1. To ensure consistency all final consensus sequences were trimmed to the WGS-t region prior to analysis.

### Analysis of public datasets

Read data from the publicly available Bioprojects was processed with MeaSeq v1.1.0 using default parameters except where noted. The additional optical duplicate removal parameter was used for BioProject PRJNA869081 as was stated in the analysis steps of the source publication [47]. Primer BED files were generated from the primers provided for each amplicon dataset to ensure amplicon regions were properly trimmed. For the Illumina amplicon dataset from PRJNA1293457, the --ivar_offset was set to 1 to assist in trimming some of the overlapping primer sites. From the sequencing date and information available for BioProject PRJNA843031 [42], we determined that “r941_prom_sup_g5014” was the most appropriate Clair3 variant calling model to use with the data. Whole genome comparisons were completed following the same method as the validation samples. Phylogenetic analysis using MeaSeq generated consensus sequences was constructed using IQ-Tree3 v 3.0.1 [52] with the general time reversible substitution matrix (GTR), gamma-distributed rate heterogeneity and 1000 bootstrapping iterations.

## Results

### Validation of MeaSeq using reference sequences

MeaSeq generated consensus sequences of surveillance targets (N450, MF-NCR) were compared to gold-standard Sanger-generated data to determine pipeline accuracy (Table 2). For the N450 region, data was available for 146 samples, and 100% concordance was achieved across all three sequencing methods. For the MF-NCR, there were 113 Sanger sequences available for comparison, and concordance values of 98.0%, 95.7%, and 95.8% for Illumina hybrid capture, Illumina tiled amplicon, and ONT tiled amplicon methods respectively. All ambiguous single nucleotide variant (SNV) discrepancies were related to the stringent MeaSeq allele frequency threshold required to call a consensus base. The ambiguous SNVs detected were all compatible with the gold-standard nucleotide but not at a high enough frequency to call a consensus at the default allele frequency (0.75 for Illumina and 0.60 for ONT). The Illumina tiled amplicon method also had one sample with a 5bp indel instead of the 6bp indel seen in the Sanger data. This sample was flagged by MeaSeq as not abiding by the rule-of-six for genome length. Upon manual inspection and curation of the read pileups in the BAM file, 84% of the reads at this position indicated an insertion of some length. We believe that the low complexity of the insertion, along with the low depth of coverage of the region, contributed to the expected 6bp insertion failing to pass the consensus threshold. As a clinical sample, this virus must abide by the rule-of-six, meaning that a 6bp insertion is the only viable insertion length. A similar insertion from another sample is depicted in Supplementary Figure 2. The ONT tiled amplicon method had one sample with one SNV nucleotide change that also related to underlying site variation seen in the sequencing reads. Due to the different wet-lab methodologies and dry-lab pipelines used and compared between studies, we cannot determine at which stage these minor differences originated.

**Table 2.** Accuracy of MeaSeq consensus sequences compared to Sanger generated N450 and MF-NCR sequences. NGS data generated by Illumina hybrid capture, Illumina tiled amplicon and ONT tiled amplicon methods.

|  | **Illumina Hybrid**<br>**Capture (n=59)** |  | **Illumina Tiled**<br>**Amplicon (n=51)** |  | **ONT Tiled Amplicon**<br>**(n=36)** |  |
| --- | --- | --- | --- | --- | --- | --- |
| **Result** | N450 (n=59) | MFNCR (n=52) | N450 (n=51) | MFNCR (n=37) | N450 (n=36) | MFNCR (n=24) |
| No. of sequences with complete consensus agreement | 59 (100%) | 50 (96.15%) | 51 (100%) | 35 (94.59%) | 36 (100%) | 23 (95.83%) |
| No. of sequences with ambiguous mismatch(es) | 0 | 2 * (3.85%) | 0 | 1 * (2.70%) | 0 | 0 |
| No. of sequences with nucleotide mismatch(es) | 0 | 0 | 0 | 0 | 0 | 1 (4.17%) |
| No. of sequences with indel(s) | 0 | 0 | 0 | 1 (2.70%) | 0 | 0 |
| Overall concordance | 100.00% | 98.00% | 100.00% | 95.70% | 100.00% | 95.83% |

\*Ambiguous bases were assigned a mismatch score of 0.5 if the ambiguity code was compatible with the reference nucleotide.

In addition to validation against Sanger generated sequences, ATCC genotype D8 and B3 reference genomes were sequenced using Illumina and ONT tiled amplicon methods and analyzed with MeaSeq using default parameters. Consensus sequences generated by MeaSeq showed 100% identity to the manufacturer-supplied reference genome sequences for both platforms.

### Characteristics of the publicly available dataset

The public dataset characteristics are summarized in Table 3 and offer a side-by-side comparison of the data obtained from NCBI and the results obtained by MeaSeq. Due to unmet inclusion criteria, nine sequences from the Illumina hybrid-capture method and 11 sequences from the Illumina tiled amplicon method were excluded from the analysis, resulting in a total of 203 sequences for comparison across the three sequencing methodologies. Genotype distribution, the amount of unique DSIds, sequences abiding by the rule-of-six and sequences with altered stop codons were identical in both datasets.

**Table 3.**
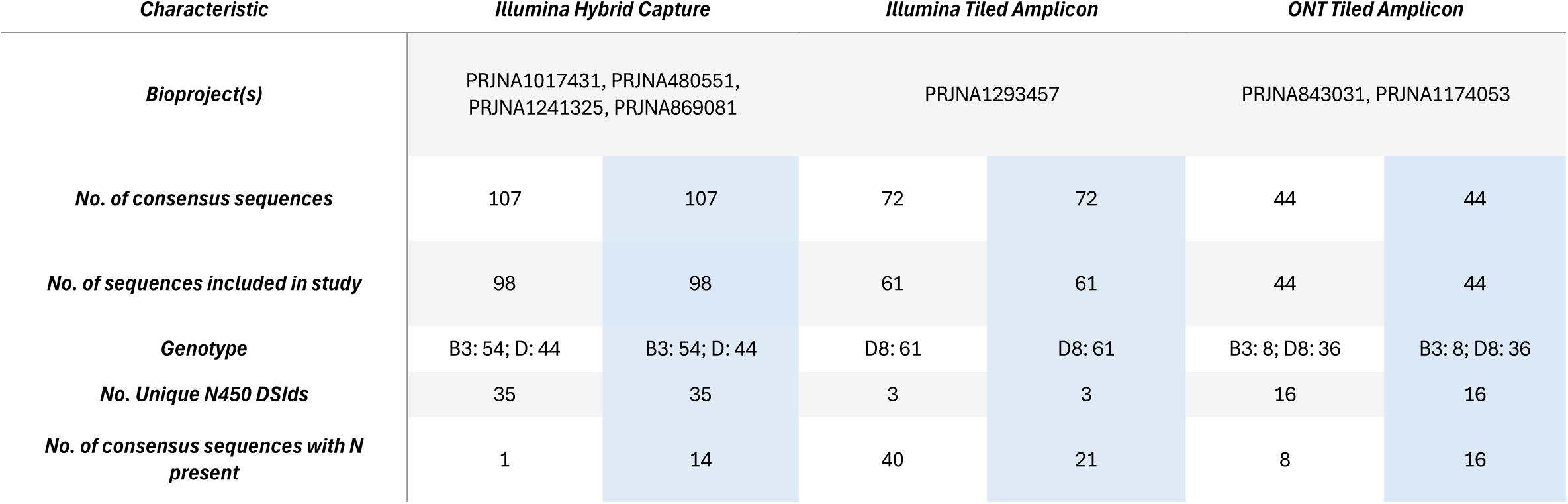
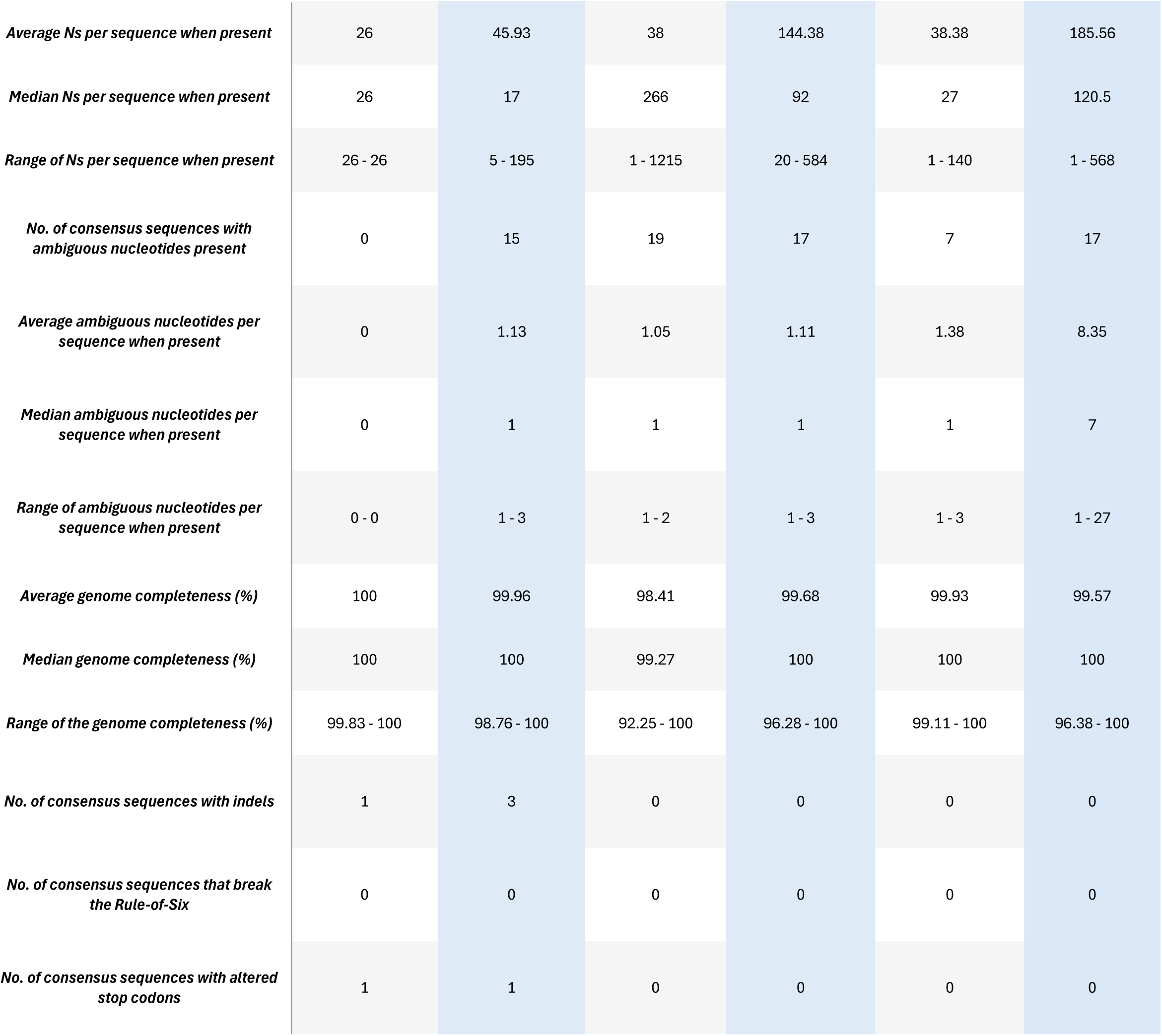
Characteristics of the publicly available dataset. A side-by-side comparison of the consensus sequences obtained from NCBI and the MeaSeq generated consensus sequences of the same read data (shaded in blue). NCBI consensus sequences may contain additional polishing or manual curation.

| Characteristic | Illumina Hybrid Capture |  | Illumina Tiled Amplicon |  | ONT Tiled Amplicon |  |
| --- | --- | --- | --- | --- | --- | --- |
| **Bioproject(s)** | PRJNA1017431, PRJNA480551, PRJNA1241325, PRJNA869081 |  | PRJNA1293457 |  | PRJNA843031, PRJNA1174053 |  |
| **No. of consensus sequences** | 107 | 107 | 72 | 72 | 44 | 44 |
| **No. of sequences included in study** | 98 | 98 | 61 | 61 | 44 | 44 |
| **Genotype** | B3: 54; D: 44 | B3: 54; D: 44 | D8: 61 | D8: 61 | B3: 8; D8: 36 | B3: 8; D8: 36 |
| **No. Unique N450 DSIDs** | 35 | 35 | 3 | 3 | 16 | 16 |
| **No. of consensus sequences with N present** | 1 | 14 | 40 | 21 | 8 | 16 |
| **Average Ns per sequence when present** | 26 | 45.93 | 38 | 144.38 | 38.38 | 185.56 |
| **Median Ns per sequence when present** | 26 | 17 | 266 | 92 | 27 | 120.5 |
| **Range of Ns per sequence when present** | 26 - 26 | 5 - 195 | 1 - 1215 | 20 - 584 | 1 - 140 | 1 - 568 |
| **No. of consensus sequences with ambiguous nucleotides present** | 0 | 15 | 19 | 17 | 7 | 17 |
| **Average ambiguous nucleotides per sequence when present** | 0 | 1.13 | 1.05 | 1.11 | 1.38 | 8.35 |
| **Median ambiguous nucleotides per sequence when present** | 0 | 1 | 1 | 1 | 1 | 7 |
| **Range of ambiguous nucleotides per sequence when present** | 0 - 0 | 1 - 3 | 1 - 2 | 1 - 3 | 1 - 3 | 1 - 27 |
| **Average genome completeness (%)** | 100 | 99.96 | 98.41 | 99.68 | 99.93 | 99.57 |
| **Median genome completeness (%)** | 100 | 100 | 99.27 | 100 | 100 | 100 |
| **Range of the genome completeness (%)** | 99.83 - 100 | 98.76 - 100 | 92.25 - 100 | 96.28 - 100 | 99.11 - 100 | 96.38 - 100 |
| **No. of consensus sequences with indels** | 1 | 3 | 0 | 0 | 0 | 0 |
| **No. of consensus sequences that break the Rule-of-Six** | 0 | 0 | 0 | 0 | 0 | 0 |
| **No. of consensus sequences with altered stop codons** | 1 | 1 | 0 | 0 | 0 | 0 |

Notable discrepancies can be found in the number of Ns and ambiguities present in the sequences. These differences can be attributed to a variety of reasons, such as biases in the different tools used to analyze the data, different parameter settings for variant calling, different primer trimming strategies, and/or manual intervention/polishing of sequences for publication. Sequences may also have been patched with targeted Sanger sequencing. Reviewing the manuscripts of the BioProjects used in this study, different settings and manual interventions were noted, which were not available for our analysis, and which likely led to higher rates of ambiguity in the MeaSeq dataset. For example, in BioProject PRJNA869081, when the depth of coverage of 10 was not met, Sanger contigs were generated to patch the consensus sequence [47]. BioProject PRJNA843031 describes that differences or ambiguities in the genome sequences were resolved manually [51]. Bioproject PRJNA1017431 tolerated no ambiguous bases and set an allele frequency threshold of 51% or greater to call nucleotides at each genomic position which differs from the Illumina MeaSeq default of ≥75% [42]. The slight genome completeness variation between datasets can likely also be explained by the different parameters and manual interventions used for each BioProject. Additionally, there were two sequences in the Illumina hybrid capture method where the NCBI data indicated a genome of 6bp shorter than the MeaSeq generated data. Primer trimming method discrepancies in the Illumina and Nanopore amplicon data also led to differences in the dataset characteristics with how primer sites and regions outside of the amplicons were handled. For an extended description of the comparison data set and for assembly statistics as well as the raw and trimmed read counts, see Supplementary Table 1.

### Diversity of Comparison samples

The diversity of the MeaSeq generated comparison dataset is presented in Figure 4. As shown, the dataset encompasses a broad range of sequences, demonstrating substantial variability across key sequence features. This diversity indicates that the dataset is well-suited for downstream analyses, as it captures a wide representation of the sequence space generated by MeaSeq. Collectively, Figure 4 highlights the depth and breadth of the dataset, supporting its robustness and suitability for this investigation.

**Figure 4.**
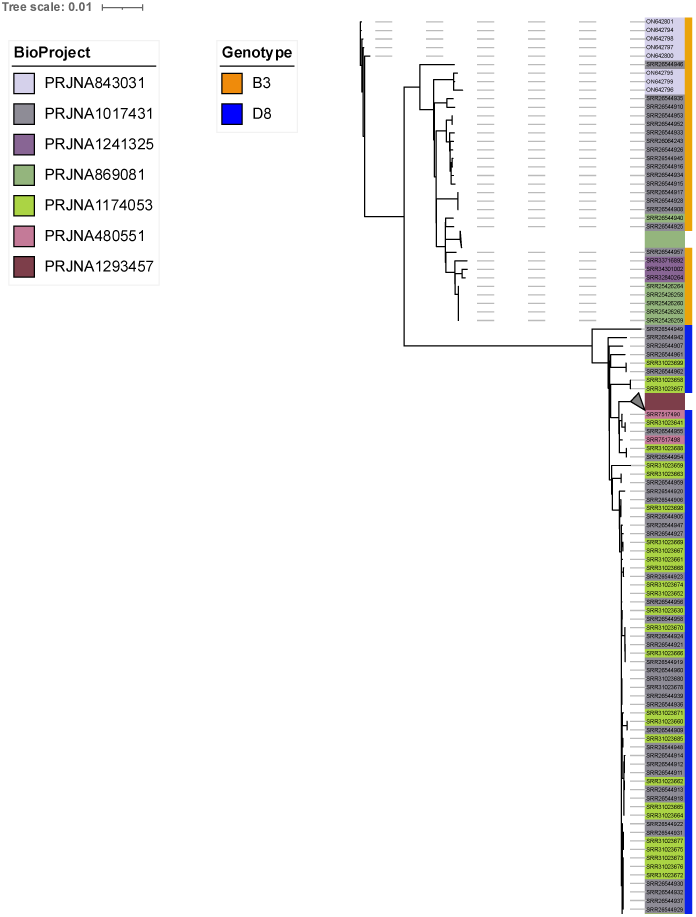
Maximum-likelihood phylogenetic tree showing the diversity of the comparison dataset. Colour annotations indicate SRA accession numbers, coloured by associated BioProject and genotype. Low diversity branches originating from the same BioProject have been collapsed.

### Evaluation of MeaSeq performance with diverse data

Performance metrics were assessed using public read data harvested from SRA that were matched to 203 consensus sequences on GenBank. The sequencing methods included three NGS strategies: Illumina hybrid capture (n=98), Illumina tiled amplicon (n=61), and ONT tiled amplicon (n=44) (Table 4). For all categorical metrics, including genotype assignment, DSId determination, rule-of-six compliance, and detection of unusual stop codons, 100% agreement was observed across all sequencing methods.

**Table 4.**
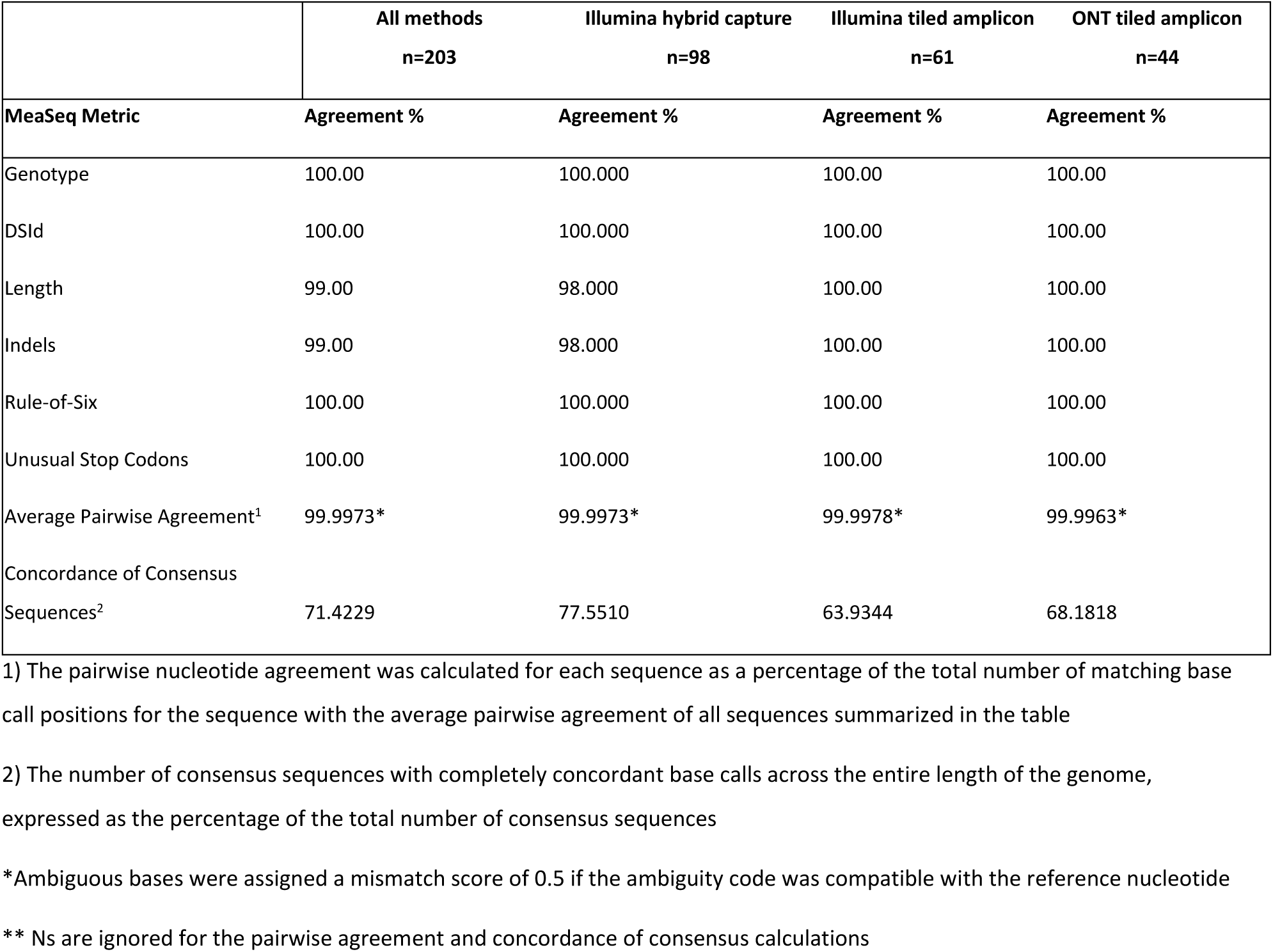
Evaluation of MeaSeq generated consensus sequences using diverse and publicly available read data matched with consensus sequences

|  | All methods<br>n=203 | Illumina hybrid capture<br>n=98 | Illumina tiled amplicon<br>n=61 | ONT tiled amplicon<br>n=44 |
| --- | --- | --- | --- | --- |
| MeaSeq Metric | Agreement % | Agreement % | Agreement % | Agreement % |
| Genotype | 100.00 | 100.000 | 100.00 | 100.00 |
| DSId | 100.00 | 100.000 | 100.00 | 100.00 |
| Length | 99.00 | 98.000 | 100.00 | 100.00 |
| Indels | 99.00 | 98.000 | 100.00 | 100.00 |
| Rule-of-Six | 100.00 | 100.000 | 100.00 | 100.00 |
| Unusual Stop Codons | 100.00 | 100.000 | 100.00 | 100.00 |
| Average Pairwise Agreement<sup>1</sup> | 99.9973* | 99.9973* | 99.9978* | 99.9963* |
| Concordance of Consensus Sequences<sup>2</sup> | 71.4229 | 77.5510 | 63.9344 | 68.1818 |

1) The pairwise nucleotide agreement was calculated for each sequence as a percentage of the total number of matching base call positions for the sequence with the average pairwise agreement of all sequences summarized in the table
2) The number of consensus sequences with completely concordant base calls across the entire length of the genome, expressed as the percentage of the total number of consensus sequences
\*Ambiguous bases were assigned a mismatch score of 0.5 if the ambiguity code was compatible with the reference nucleotide
\*\* Ns are ignored for the pairwise agreement and concordance of consensus calculations

There were two sequences, both belonging to BioProject PRJNA1241325 that differed in length due to indels. This was due to a well-supported, low complexity 6bp insertion identified in a region with a strong negative strand bias in all hybrid capture samples (Supplementary Figure 2). For all methods, the average pairwise agreement achieved by MeaSeq compared to the published consensus sequence was 99.9973%. Separated by method, the average pairwise agreement was 99.9973%, 99.9978%, and 99.9963% for Illumina hybrid capture, Illumina tiled amplicon, and ONT tiled amplicon, respectively.

Across all three methods, 71.42% of MeaSeq generated sequences were in 100% concordance with the publicly available comparative data. Minor disagreements observed in the average pairwise agreement between the public data and the MeaSeq generated data occurred at the individual nucleotide level and could be attributed to site-specific ambiguity, primer trimming of amplicon data, and the differing variant calling thresholds used for calling a consensus or an ambiguous base by the different analysis workflows. In addition to these thresholds, publications associated with BioProjects PRJNA869081, PRJNA1017431 and PRJNA843031 also mention manual intervention in some of their consensus sequences which, although a common practice, cannot be replicated from the raw data alone [42, 47, 51]. Primer trimming tool choice also plays a role in the underlying nucleotide discordance differences seen in primer regions where the complicated amplicon scheme reads may not be fully primer trimmed leading to ambiguous or masked sites (Supplementary Figure 1). We did not encounter any nucleotide disagreements that could not be explained by these methodological factors.

Repeatability was also assessed by running the MeaSeq workflow under the same conditions twice on a random subset of samples (n=20) for each of the three sequencing methods. For all methods, after repetition, the exact same results were achieved 100% of the time (n=60) showing that MeaSeq produces repeatable outputs.

## Discussion

NGS is transforming MeV surveillance. As historic vaccination successes have reduced circulating MeV diversity to only two strains, the discriminatory power of the established N450 surveillance region is diminished. WGS data offers better tracking of transmission chains and resolution of outbreaks. With declining costs of NGS and more readily available library preparation protocols, WGS is becoming more widely adopted in the surveillance of infectious diseases [53]. However, not all scientists are trained in bioinformatics, and this can lead to a bottleneck in the analysis and interpretation of NGS data. With specific genomic challenges and reporting requirements, a bioinformatics pipeline specific for MeV that is accurate, easy to use, automated, and robust would enhance the measles scientific community and their surveillance efforts.

Here we have presented MeaSeq, a bioinformatics pipeline developed for direct analysis of MeV whole genome sequence reads obtained with Illumina or ONT platforms. This pipeline incorporates established viral analysis workflows for variant detection and consensus building with additional capabilities designed specifically for operational MeV analysis and reporting. These focused analyses include MeV specific genotype and DSId assignment, quality control checks such as genome divisibility and gene validation, subconsensus nucleotide analysis and mixed-site highlighting, and genomic plotting. All outputs are summarized in a final report that allows users to analyze, interact with, and share their data.

The MeaSeq pipeline was first validated against reference sequences using NGS read data generated from a variety of sequencing approaches with minimal, low impact differences seen between the MeaSeq-consensus sequences and the reference sequences. All MeaSeq generated sequences completely matched the Sanger-generated N450 genotyping region and the ATCC reference genomes demonstrating the high accuracy of the MeaSeq pipeline. Minimal differences were seen in a minority of samples when compared to the Sanger-generated MF-NCR region. As the WGS methods required different wet lab procedures from the Sanger reference sequences, we cannot determine explicitly if the small subset of differences arose from the bioinformatic analysis, the wet lab method, or a combination of both.

The Illumina amplicon tiled dataset included one sample with an indel that was not detected in the Sanger data. The rule-of-six divisibility check built into MeaSeq flagged this sample as not divisible so that it could be manually inspected and resequenced if needed. In some cases, MeaSeq identified sites that did not meet the threshold of a major allele call where the corresponding Sanger data did not indicate support for multiple alleles. NGS has an advantage over Sanger where these mixed sites can be identified and more thoroughly examined to explore subconsensus nucleotides and potential viral evolution. These sites had underlying mixtures that were traced back to the reads. The MeaSeq default to call a major allele frequency is set to 0.75 for Illumina and 0.60 for ONT data and in these cases, the pipeline is behaving as expected. A deeper investigation into these subconsensus nucleotides may aid outbreak and transmission chain investigations or assist in detecting potential sequencing errors in the dataset. A custom figure produced by MeaSeq identifying positions with alternate alleles present in the pileup above 15% simplifies this process (Figure 3A). The stringency of the default parameters being responsible for a slight increase in ambiguity when the data quality (depth, complexity) is lower, which can be common for the MF-NCR region.

The MF-NCR is of interest in MeV analysis as its hypervariability has been identified as a potential surveillance target [6]. The region is known to be difficult to sequence, as shown by Supplemental Figure 3, where there is a noticeable drop in depth and quality across the samples. The MF-NCR contains many homopolymer stretches, which are challenging for most polymerases to sequence [54], and is also GC-rich making it prone to secondary structures [55]. Bioinformatically, it is hard to assemble due to the genomic repeats and homopolymer structures that create challenges for basecalling and read alignment in the region [56]. As a non-coding region, it is not transcribed for translation into protein and is consequently less represented than protein-coding regions. These non-coding regions are more tolerant of indels as they will not cause any disruptions to a protein sequence that would be detrimental to virus survival. As a result, the analysis of the MF-NCR needs to be done with caution ensuring that the coverage of the region is sufficient for accurate basecalls. These MeV-specific genomic constraints have been considered in the development of MeaSeq and the optimization of suitable variant thresholds, providing an advantage over generic viral WGS pipelines.

Evaluation with the publicly available dataset provided near perfect sequence matches producing an average percent identity of 99.9973%. The minor differences between bioinformatic analyses can be attributed to the sequencing depth, breadth, and decisions on the thresholds used for variant calls. Areas of low depth in the MF-NCR were the largest cause of discordance between analyses with ambiguous positions and nucleotide mismatches stemming from differing variants quality and allele frequency thresholds along with manual curation from sequence submitters [42, 47, 51].

Amplicon primer regions were also identified as a cause of ambiguous bases and mismatches highlighting the need for robust primer mapping and thorough primer trimming in tiled amplicon datasets. Tiled amplicon sequencing is widely adopted for viral sequencing analysis with primer schemes available for many different pathogens. This approach involves designing primers accounting for genomic variability and amplifying the multiplex in two or more pools to create amplicons that slightly overlap each other without creating short, fragmented products [57]. This method helps in sequencing samples with lower cycle thresholds (Ct), tough to sequence regions, or with viruses that have a lower viral copy number in clinical samples. In the comparison dataset, the largest amount of ambiguity and mismatches occurred in the primer regions of tiled amplicon sequences. This suggests that proper primer trimming may be inadequate with some of the wetlab assay designs, leading to the introduction of sequence artefacts.

Quality control checks are essential for analyzing and releasing MeV whole genome data as MeV has strict biological constraints for viability including the rule-of-six and gene validity [3]. The ability to flag critical biological inaccuracies without removing the sequence from downstream reporting is especially helpful for dealing with samples that range in quality, such as those included in our comparison dataset. The quality warnings and visualizations included in the MeaSeq report will assist users with minimal bioinformatics experience in the interpretation of their whole genome data and in any potential manual sequence curation that is required.

Gene functionality is an important biological aspect to track in the analysis of MeV genomes. The MeaSeq pipeline identifies any frameshift mutations and stop codon variants in coding sequences that could affect the virus’ viability. In the current analysis, the MeaSeq report was able to correctly identify known anomalies such as the late stop in the Phosphoprotein of SRR2606424 (Figure 3A) and an indel event in the P/V/C gene from BioProject PRJNA1017431. In both events, the sample warning flag prompts the user to investigate further and confirm the result. In the case of the late Phosphoprotein stop codon, further analysis showed the new stop located downstream by 19 amino acids [5].

This study has a limitation. Ideally, validation would be performed using a well-characterized, genetically diverse reference panel comprising samples from multiple sources and generated under standardized conditions. In the absence of such a resource, our analysis was necessarily limited to Sanger-generated sub-genomic fragments and publicly available datasets with raw sequencing reads deposited in the NCBI SRA, which varied in sequence quality and metadata completeness. Despite the limitations of the publicly available data, the results support the utility of MeaSeq across a diverse set of real-world data.

The measles genomics community would greatly benefit from a coordinated effort to compile and curate a comprehensive reference dataset suitable for method validation and benchmarking.

Nonetheless, MeaSeq was evaluated using a phylogenetically diverse collection of samples derived from multiple sources and incorporating three different next-generation sequencing methodologies.

Importantly, the dataset included sequences containing insertions and deletions, altered stop codons, and a range of sequence qualities. As such, the validation panel represented a challenging and realistic test environment, providing a robust assessment of MeaSeq’s performance as a bioinformatics pipeline.

In summary, we have developed and thoroughly validated MeaSeq, a bioinformatic pipeline for the analysis of MeV genomes generated from short and long read NGS platforms. With the strict biological constraints of the MeV genome, an increasing worldwide case count, and the decreasing genomic diversity in the standard N450 genotyping region, a well validated MeV specific NGS pipeline has a key place in assisting MeV researchers and public health scientists. This rapid and easy-to-use pipeline will enhance NGS analysis and surveillance in the MeV scientific community where such a pipeline does not currently exist. It has been validated on numerous surveillance samples of varying degrees of quality to highlight functionality and accuracy and provides an interactive report serving as a platform for the user to further explore their data. Future directions aim towards expanding the outputs available for researchers and implementing feedback from the broader measles research community.

## Author Contributions

Conceptualization: D.H., V.Z., A.T.D., J.H.; Data curation: D.H., A.A., V.Z.; Formal analysis: D.H., A.A., V.Z.; Investigation: S.V.D., S.A., V.Z.; Methodology: D.H., A.A., V.Z., A.T.D.; Supervision: A.T.D., J.H.; Validation: D.H., V.Z.; Visualization: D.H., V.Z., A.A., M.P.; Writing – original draft: D.H., V.Z., A.A., M.P.; Writing – review and editing: all authors.

## Funding sources

This study was supported by the operating budgets of the Computational and Operational Genomics

Section and the Measles, Mumps, and Rubella unit at the National Microbiology Laboratory Branch.

## Acknowledgements

We gratefully acknowledge our provincial partners who refer specimens to the NML for measles sequence surveillance. We would also like to acknowledge the Genomics Core facility at the NML for graciously running our products for the Sanger and Illumina Sequencing and James Robertson for helpful comments on the manuscript draft.

