## Supplementary Figures for "Evaluation of MeaSeq: comprehensive analysis and reporting of measles virus whole genome sequences"

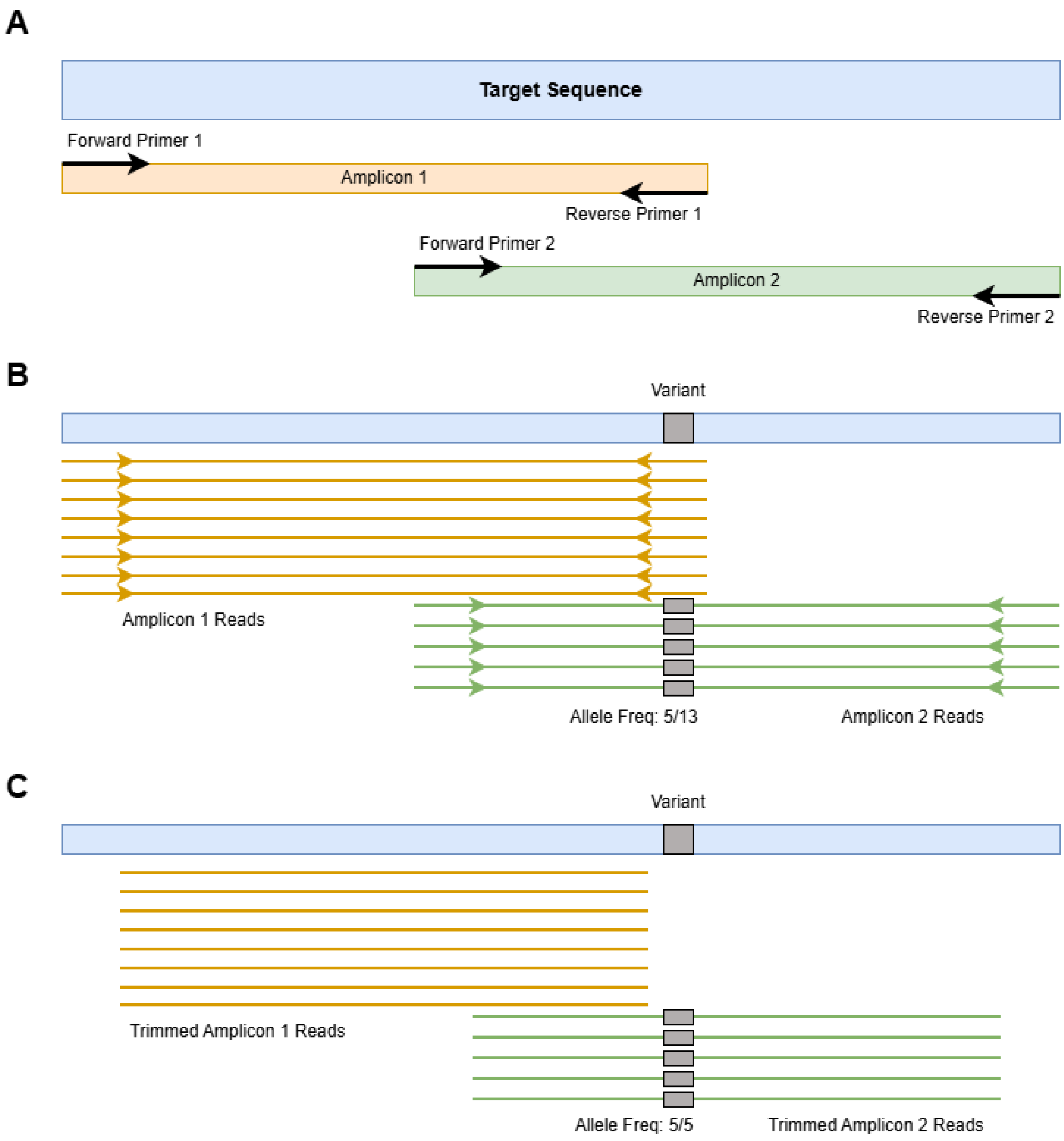


**Supplemental Figure 1.** Schematic diagram of how primer trimming affects variant calls. A) Generic two-pool amplicon schema for the target sequence. B) Visualized pileup of the sequencing reads for the target sequence, displaying how the variant in the target may be masked without proper primer trimming. C) The same visualized pileup of the sequencing reads with the reads correctly primer trimmed showing the allele correctly called.

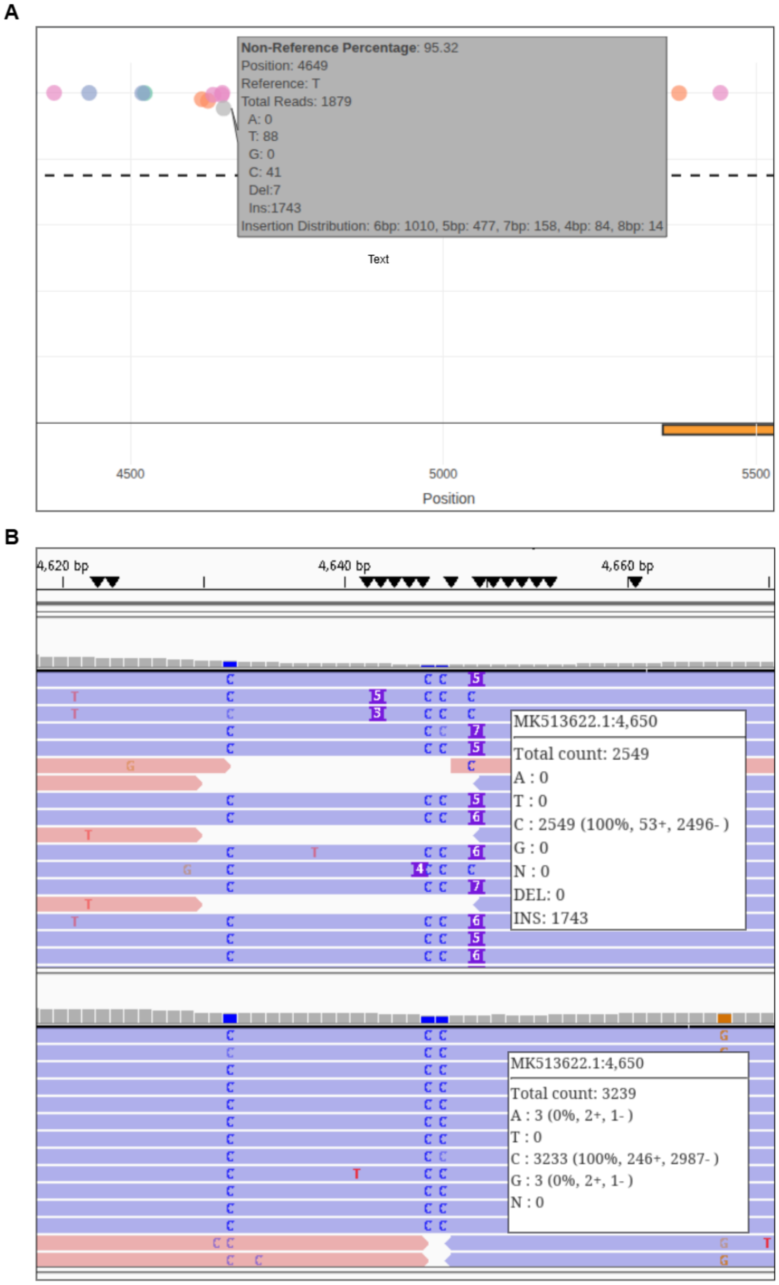


**Supplemental Figure 2.** Insertion site pileup summaries from MeaSeq and IGV. A) The MeaSeq subconsensus variant plot shows the distribution of insertion calls underlying the site. B) IGV shows the strand bias and insertion calls for a sample with the insertion (top) and without (bottom). Overall, there is a consensus to call an insertion of 6bp. The distribution of insertion lengths stems from the low-complexity of the insertion with a 6bp insertion making the most sense due to the rule-of-six genome organization. Negative strand bias at the site is seen in samples with and without the insertion.

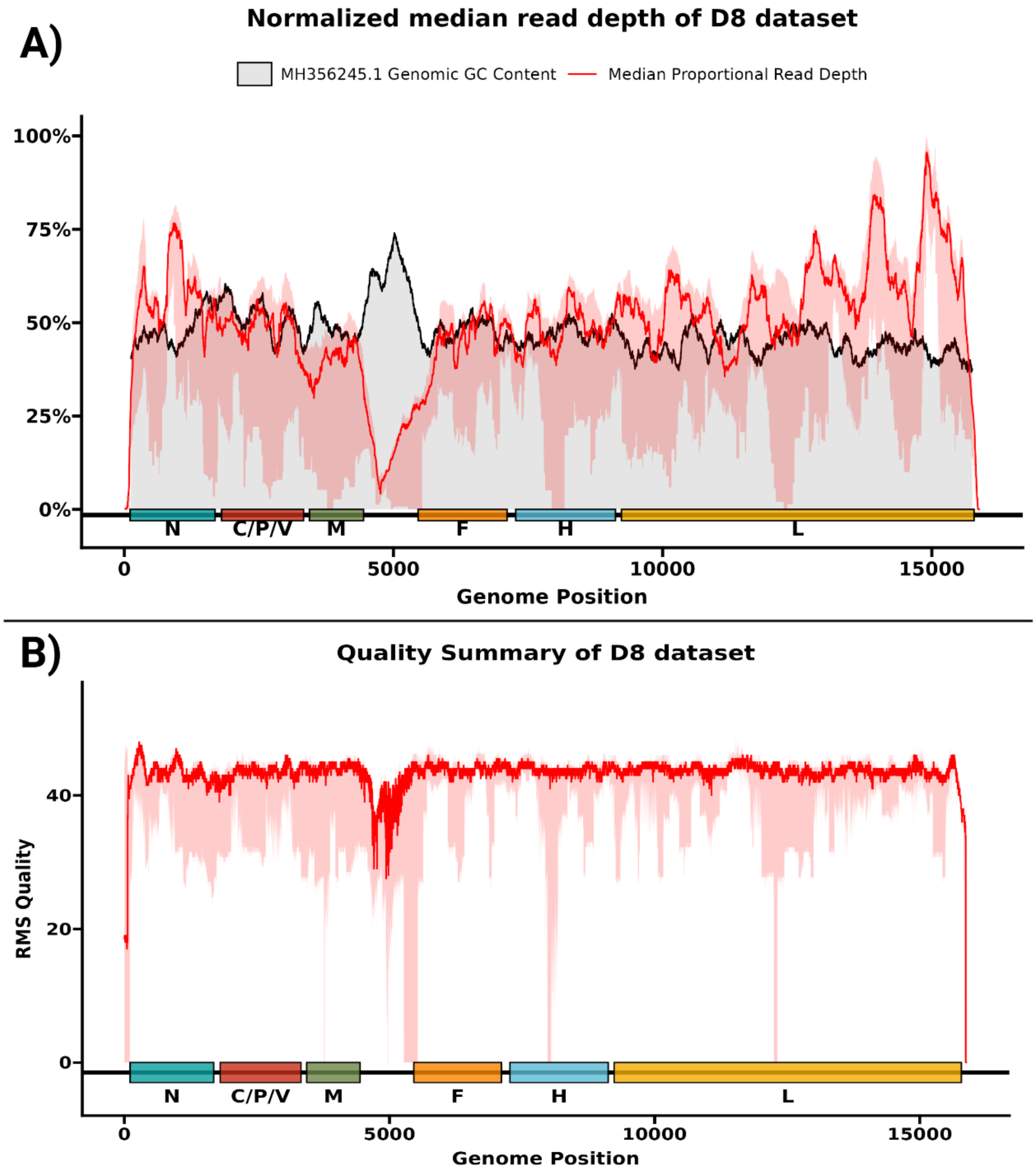


**Supplemental Figure 3.** Whole genome read depth and quality in D8 hybrid capture data. Median read depth across the D8 MeV genome, normalized to the proportion of the maximum positional depth per sample, is shown in A (red). Reduced depth in the MF-NCR is co-located with an area of GC richness in the D8 reference genome (black/grey). Median root-mean squared (RMS) base quality is shown in B. In both plots, the lighter red shaded area indicates the 0.25-0.75 IQR. MeV gene annotations along the x-axes are based on D8 reference MH356245.1.
